# *ERas-*Null Mice Generated Directly from Embryonic Stem Cells in a Lipid-rich Medium Enable the Discovery of A Novel Role in Craniofacial Development

**DOI:** 10.1101/2023.11.01.565209

**Authors:** Yiren Qin, Qiyu Tian, Hoyoung Chung, Fuqian Geng, Daylon J. James, Jianlong Wang, Duancheng Wen

## Abstract

Generating mice entirely from embryonic stem cells (ESCs) via tetraploid complementation, termed all-ESC mice, is considered the gold standard for pluripotency testing. In this study, we examined the effects of our newly reported lipid-rich 2i/LIF medium (2iLA) on ESCs for genetic engineering. As a proof of principle, we targeted the *ERas* gene using the 2iLA medium, since *ERas*-null mice displayed no noticeable phenotype in a previously developed mouse model through chimeras. We demonstrated that the 2iLA medium effectively supports the creation of both male and female *ERas*-null all-ESC mice, emphasizing the advantages of this culture medium. Unexpectedly, we observed that *ERas*-null all-ESC mice produced with the 2iLA medium exhibited a non-Mendelian lethal craniofacial anomaly, which can be mitigated by using the lipid-free 2i/LIF medium. Our findings not only highlight the potential of the 2iLA medium for gene targeting but also reveal a novel lipid-associated role of the *ERas* gene in craniofacial development. Our system offers a unique alternative for studying developmental gene functions unachievable with traditional methods and provides a novel platform for the rapid construction of mouse models.

**Graphic Abstract:** All-ESC pups from wild-type (WT) ESCs cultured in lipid-rich 2iLA medium are normal. In contrast, both male and female *ERas*-null isogenic all-ESC pups produced from ESCs in 2iLA medium display a non-Mendelian craniofacial anomaly. This phenotype is alleviated when the ESCs are grown in a lipid-free 2iL medium.

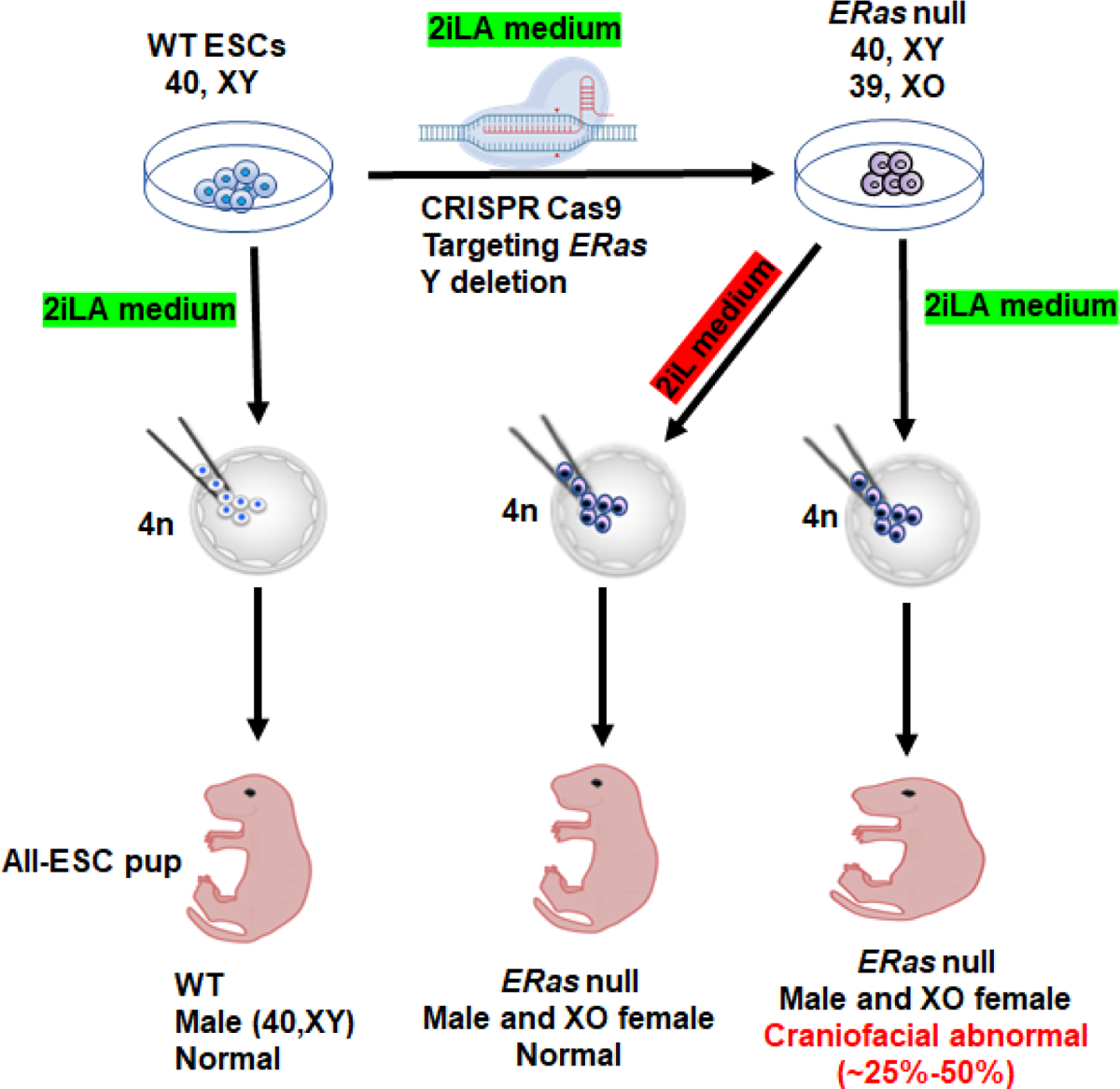

## Introduction

Genetically modified (GM) animals are essential tools for studying fundamental biology and human diseases. A typical approach for creation of GM mice involves the generation of tetra-parental chimeras from normal embryos and GM embryonic stem cells (ESCs), followed by multiple rounds of breeding to obtain both male and female homozygotes for germline propagation of the mutations. However, this process is generally time-consuming, laborious and costly (1).

Mouse ESCs, derived from the inner cell mass of blastocysts, possess unlimited self-renewal and differentiation capacity if maintained in their ground-state pluripotency (2–4). Pure ESC-derived mice (all-ESC mice) can be directly and efficiently generated through tetraploid complementation (referred to as 4n competency), a process involving the injection of ground-state ESCs into tetraploid (4n) blastocysts. In these 2n/4n chimeric embryos, 2n ESCs contribute to the entire embryo proper, while the host 4n cells can only form the extra-embryonic tissues during embryogenesis (5–7). We have recently demonstrated that isogenic male and female all-ESC mice can be created through targeting the Y chromosome in XY male ESCs using CRISPR technology and tetraploid complementation (1).

Initially, pluripotent mouse ESCs were established and cultured in medium supplemented with fetal bovine serum on feeders. However, the presence of undefined serum components led to heterogeneous cultures, resulting in a gradual loss of pluripotency (8) and the 4n competency. The breakthrough discovery that inhibition of Mek1/2 and Gsk3β (2i) maintains ESCs in a more homogeneous state of naive pluripotency (4) in the medium supplemented with leukemia inhibitory factor (2i/LIF).

However, prolonged culture of mouse ESCs in 2i/LIF unfortunately leads to aneuploidy, DNA hypomethylation, and loss of imprinting, which impairs the 4n competency (9–11). We recently reported that supplementing lipid-rich albumin (AlbuMAX) into 2i/LIF medium (2iLA) can significantly improve the genomic stability and 4n competency of mouse ESCs over long-term culture *in vitro* (11). Our innovative culture medium, combined with the technology to generate isogenic male and female all-ESC mice, provides a feasible approach to expedite the generation of GM mouse model from ESCs.

*ERas* encodes a Ras-like GTPase protein first identified as an embryonic stem cell-specific Ras variant (12). Recent studies suggest that *ERas* could be a potential marker for formative pluripotency (11, 13–15), and its expression is notably elevated in the E5.5–7.5 endoderm and epiblast (16). *ERas* has been demonstrated to have a significant impact on ESC proliferation and to enhance somatic cell reprogramming (17, 18). Although *ERas* is highly expressed in early mouse peri-implantation embryos (16), it is not crucial for either pluripotency or embryonic development (12). The exact role of *ERas* gene in early development remains unclear.

In this study, we assessed the impact of 2iLA medium on *in vitro* genetic manipulations of ESCs by targeting the *ERas* gene, known to be non-essential for pluripotency, and by deleting the Y chromosome to generate isogeneic 39, XO female ESCs. We found that bona fide pluripotency was preserved in 2iLA medium following the extensive genetic engineering of ESCs, as demonstrated by the effective production of both male and female *ERas*-null all-ESC mice via tetraploid complementation.

Surprisingly, we also observed that *ERas*-null all-ESC pups display a non-Mendelian craniofacial abnormality specifically from cells cultured in the lipid-rich 2iLA medium. Our results reveal a previously unidentified role of the *ERas* gene in directing craniofacial development, potentially linked to lipids during early development.

## Results

### The lipid-rich 2iLA medium supports extensive genetic manipulations in ESCs while preserving bone fide pluripotency

Supplementing the standard 2i/LIF medium with lipid-rich albumin, AlbuMAX, enhances long-term genome stability while promoting a transition in pluripotency to a formative-like state (11). In line with previous study, the *ERas* gene exhibits a notable increase in expression in formative-like ESCs cultured in the 2iLA medium (**Fig. 1A**), a finding further confirmed by western blotting analysis at the protein level (**Fig. 1B**).

**Figure 1.**
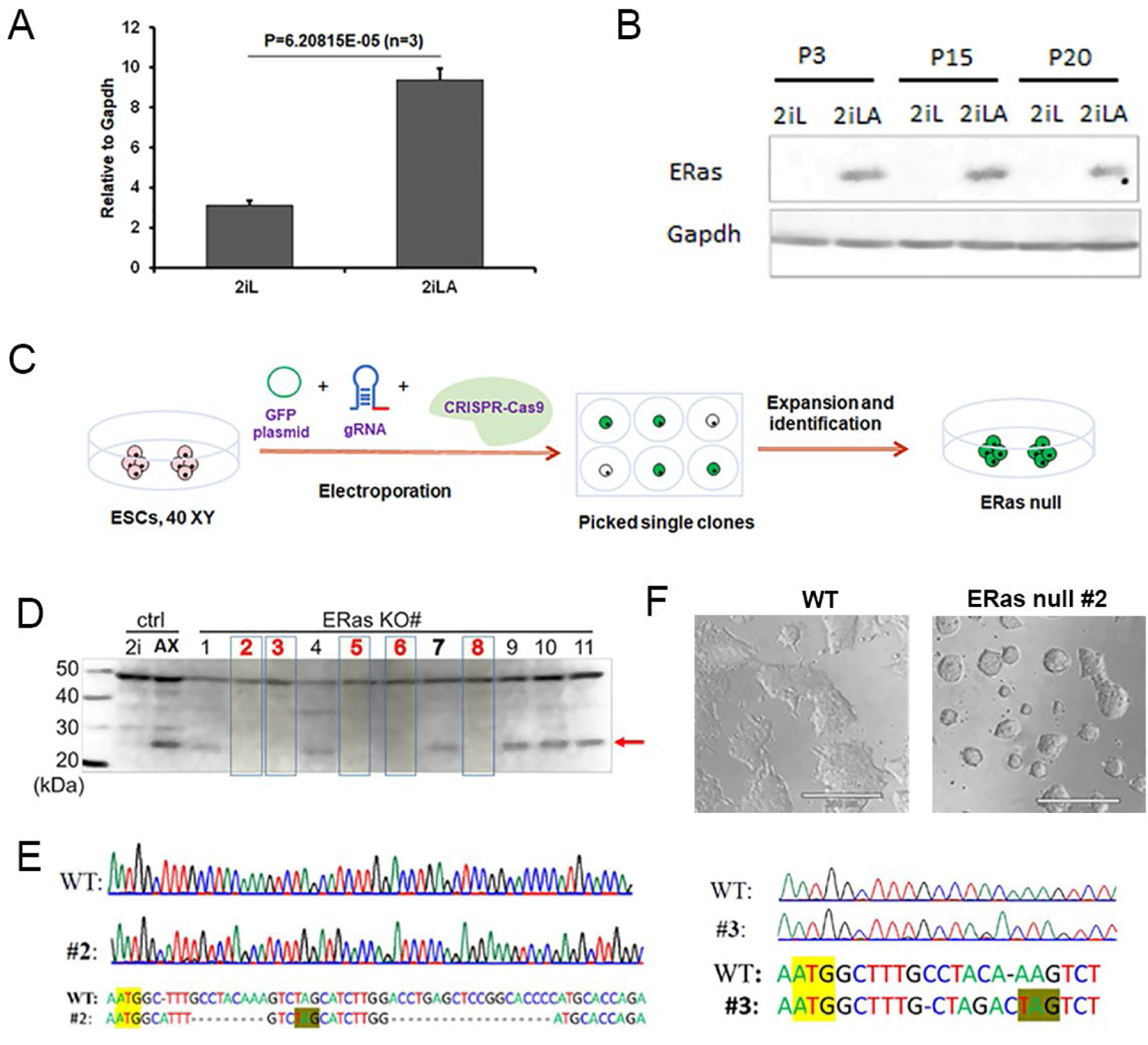
Generation of *ERas-*null ESCs via CRISPR/Cas9 in 2iLA medium. A) Real-time PCR analysis of ERas expression in ESCs cultured in 2i/LIF (2iL) and AlbuMAX-supplemented 2i/LIF (2iLA) medium. B) Western blot analysis shows the absence of ERas protein in ESCs cultured in 2iL but presents in ESCs cultured in 2iLA medium. C) Schematic representation of the ERas knockout strategy using CRISPR/Cas9. D) Western blot analysis reveals the absence of ERas protein in the subclones (the arrow points to the expected ERas band) after CRISPR targeting. E) Sequencing results for ERas-null ESC clones #2 and #3. F) Contrasts in colony morphology between ERas-null ESCs, which exhibit a domed naïve morphology (right), and wild-type ESCs, which display a flat colony appearance (left) when cultured in 2iLA medium.

The *ERas* gene is located on the X chromosome and male ESCs possess only one X chromosome. We designed guide RNAs to target the first exon using the Guide Design Resource from the Zhang lab (http://crispr.mit.edu/). We utilized purified Cas9 proteins with nuclear localization signals that can form functional gRNA-Cas9 ribonucleoprotein complexes (RNPs) *in vitro* and introduced the RNPs into 2iLA-cultured ESCs via electroporation (**Fig. 1C**). Approximately 48 hours after electroporation, the ESC colonies were digested with trypsin, and single GFP+ cells were manually isolated using a micromanipulator under fluorescent microscopy. These GFP+ cells were individually seeded into 96-well plates on feeder layers for clonal expansion in 2iLA medium. Typically, cell colonies emerged after one week in culture and were then transferred to gelatin-coated dishes for 1-2 passages in 2iLA medium before genotyping (**Fig. 1C**).

From the single-cell GFP+ cells, we derived 15 clones. Of these, five did not express the *ERas* protein, as determined by a western blotting assay cultured in 2iLA medium (**Fig. 1D**). Sequencing these five clones revealed that the targeted *ERas* gene had excisions with either indels or small insertions (**Fig. 1E, Supplemental File 1**).

These alterations caused frame shifts, resulting in the successful depletion of the *ERas* protein and loss of function in these clones. The sequencing analysis of potential off-target sites revealed that there were no off-target indels in the targeted clones (data not shown). Notably, the *ERas*-null ESCs cultured in 2iLA medium exhibited a domed naïve colony morphology, whereas the WT ESCs displayed an expected flat and large formative colony morphology (**Fig. 1F**). This observation confirms the loss of function of the *ERas* gene, which is believed to play a role in the transition to formative pluripotency.

We targeted the ESC line with proved capable of generating all-ESC mice through tetraploid complementation. To determine if ESCs cultured in 2iLA medium still retain the potential to produce all-ESC mice after genetic editing using CRISPR, we tested the five *ERas*-null ESC lines with tetraploid complementation, aiming to generate all-ESC mice (**Fig. 2A**). Out of these five *ERas*-null ESC lines, three successfully produced all-ESC pups, with a success rate ranging from 10-39% of the transferred embryos (**Fig. 2B**). This efficiency is comparable with that of the WT parental ESCs (11), indicating that the 2iLA medium effectively preserves the pluripotency of ESCs even after genetic modifications with CRISPR.

**Figure 2.**
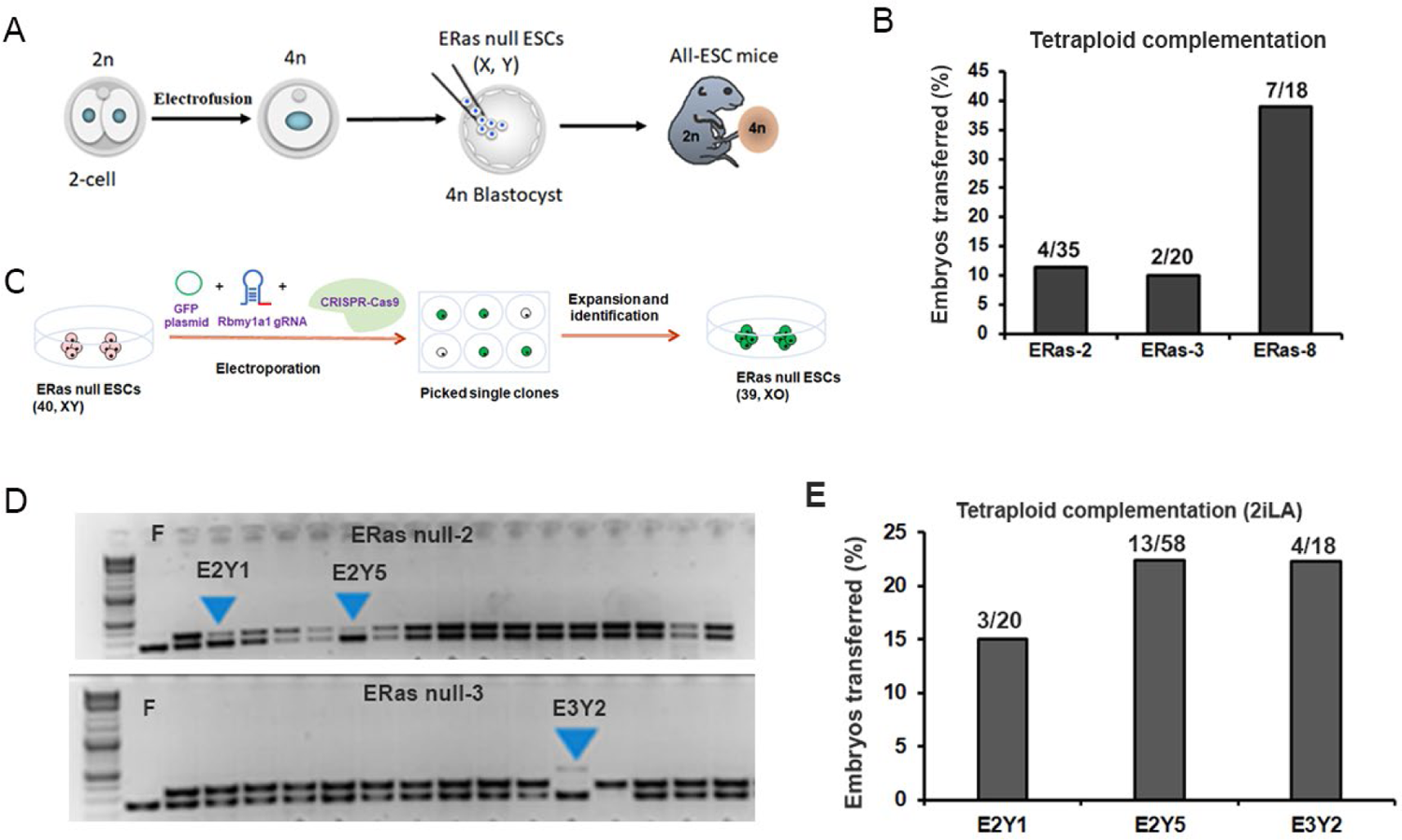
Generation of both male and female *ERas*-null all-ESC male mice via tetraploid complementation. A) Illustration depicting the process for producing ERas null male all-ESC mice through tetraploid complementation. B) Efficiency of generating all-ESC male mice using ERas null male ESCs. Numbers on each bar indicate the number of pups/number of embryos transferred. C) Schematic representation of the generation of XO female ERas-null ESCs via Y chromosome deletion. D) Genotyping of the Uba1y gene in the targeted GFP+ subclones. The arrow points to the deletion of the Y chromosome. F: female control. E) Efficiency of XO ERas-null ESCs in generating all-ESC pups via tetraploid complementation.

We also generated monosomic female XO all-ESC mice by deleting the Y chromosome in male ESCs. This was achieved by targeting the Y chromosome-specific *Rbmy1a1* gene, a technique we previously developed in our laboratory (1). To assess if the 2iLA medium can support consecutive rounds of genetic manipulation, we further targeted the *ERas*-null clones for Y chromosome deletion. Briefly, we introduced the Cas9 protein, gRNA, and GFP reporter plasmid into the *ERas*-null ESC lines via electroporation (**Fig. 2C**). Once the transfected cells were seeded and the GFP+ single-cell clones expanded, genotyping was performed to verify the status of the Y chromosome. From this process, we identified three subclones that were missing the Y chromosome specific *Uba1y* gene (**Fig. 2D**), indicating the loss of Y chromosome in the ESCs. These XO *ERas*-null ESCs were then subjected to tetraploid complementation to produce all-ESC pups. Remarkably, the three subclones produced all-ESC pups with efficiencies ranging from 15-22% (**Fig. 2E**). These results demonstrate the maintenance of pluripotency of these XO *ERas*-null ESCs even after two rounds of genetic manipulations, highlighting the robust support for *in vitro* ESC targeting using the 2iLA medium.

### *ERas*-null all-ESC pups display a non-Mendelian craniofacial abnormalities

Remarkably, among the *ERas*-null all-ESC male pups, a significant proportion (ranging from 25% to 50%) display pronounced craniofacial abnormalities (**Fig. 3A, 3B**), manifesting as a non-Mendelian phenotype. Those pups with these pronounced defects fail to survive long after birth. On the other hand, those without these deformities develop normally and grow into fertile adults. Given that these consistent non-Mendelian craniofacial developmental anomalies were observed across three different ESC clones, each with unique indels in the *ERas* gene, and since other pups remain unaffected, we infer that this observed phenotype is more likely a result of the loss of *ERas* function than potential off-target effects. Notably, the monosomic XO ERas-null all-ESC pups exhibited the same non-Mendelian craniofacial abnormalities (**Fig. 3C, 3D**), suggesting that this phenotype is consistent across both sexes.

**Figure 3.**
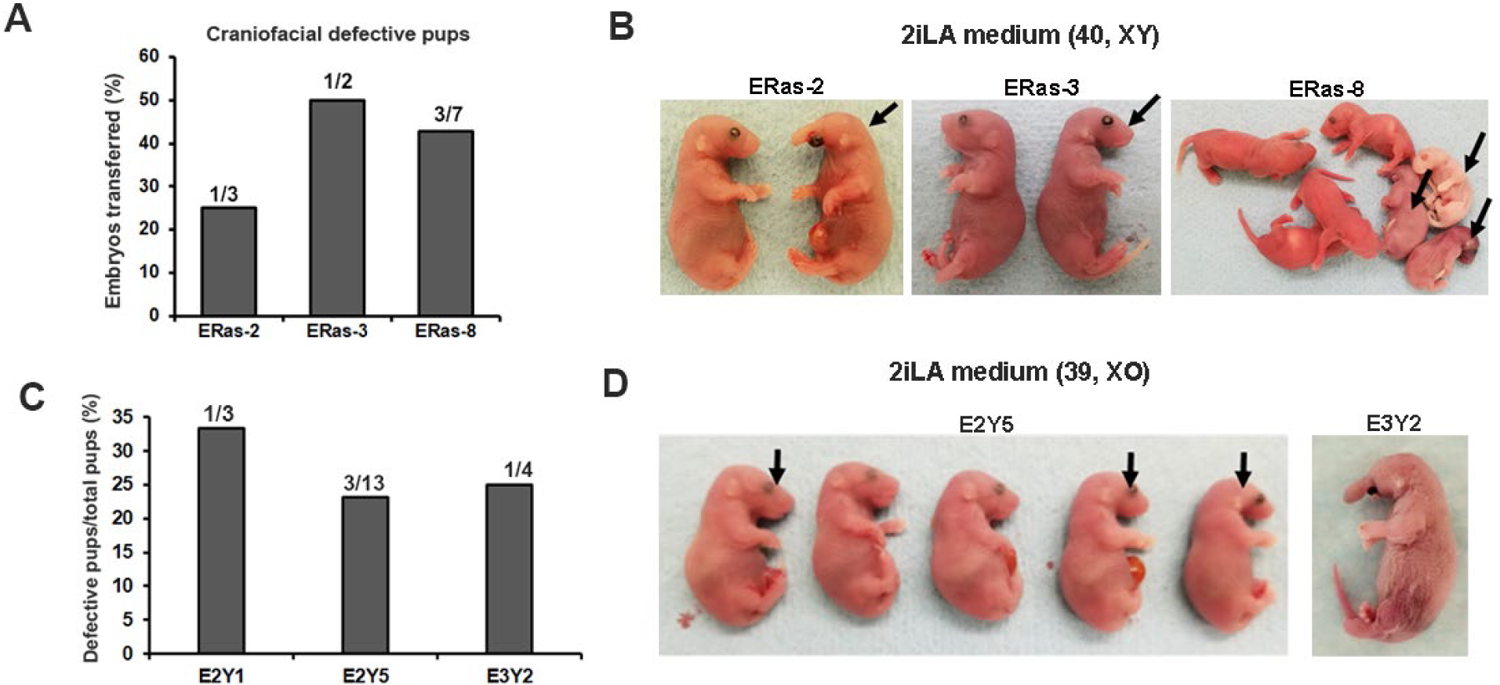
Both male and female *ERas*-null all-ESC mice display a non-Mendelian craniofacial phenotype. A) Frequency of abnormal pups observed in *ERas*-null all-ESC mice. The numbers on each bar represent the defective pups/total pups. B) Congenital anomalies identified in the *ERas*-null all-ESC pups derived from three specific ESC clones (#2, #3, #8). Pups with defects succumb shortly after birth, whereas other pups appear normal. C) Frequency of abnormal *ERas*-null monosomic XO all-ESC pups cultivated in 2iLA medium. D) Developmental abnormalities observed in XO female *ERas*-null all-ESC pups originating from subclones E2Y5 (left) and E3Y2 (right).

Histological analyses of *ERas* null all-ESC pups with pronounced defects revealed a spectrum of congenital craniofacial malformations that were incompatible with sustaining life. Common malformations observed in all pups included varying degrees of holoprosencephaly (HPE), microstomia with either mandibular hypoplasia or agnathia (absence of the mandible), micro- or aglossia (rudimentary or absent tongue), and open eyelids, which were associated with potential eyelid hypoplasia and corneal necrosis (**Fig. 4A-4D**). Among the six pups examined, three exhibited severe malformations in addition to the common anomalies mentioned above. These severe malformations included synophthalmia, arrhinia with frontal proboscis formation, central incisor or adontia, synotia, alobular holoprosencephaly, and possible agenesis of abdominal organs (**Fig.4A-4D, Supplemental File 2**).

**Figure 4.**
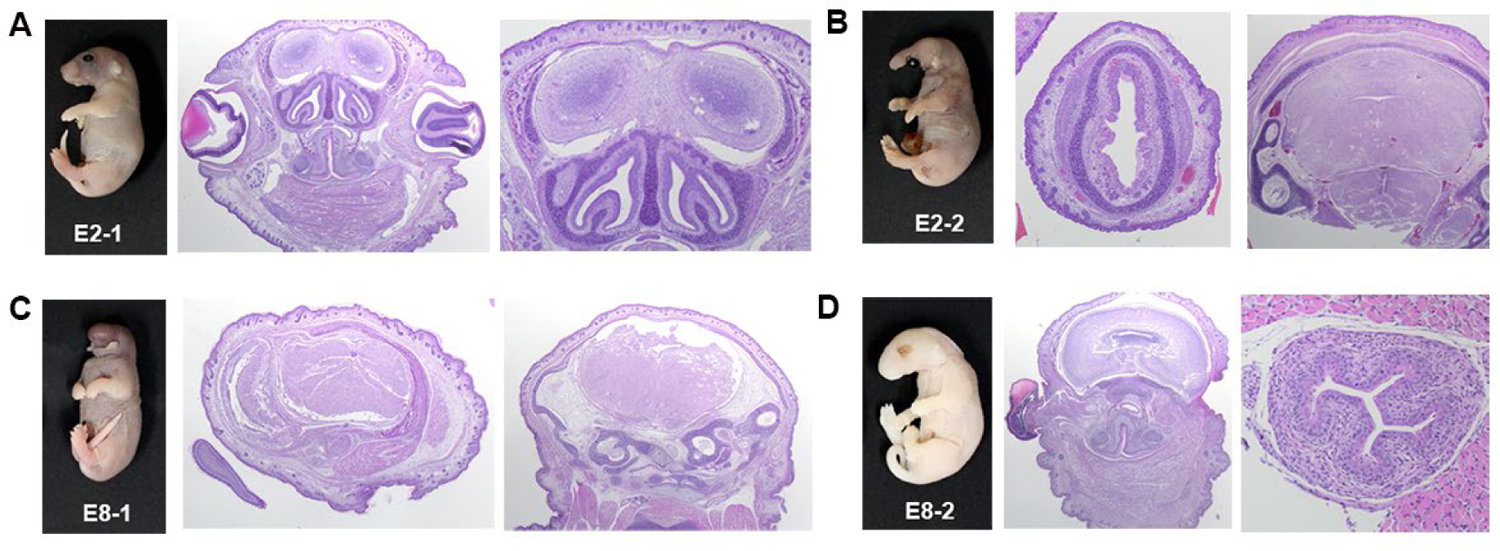
Pathological analysis of *ERas*-null all-ESC pups (P0). A) The E2-1 all-ESC pup derived from ERas null ESC clone-2 exhibited well-developed head, body, and extremities, but displayed Microstoma with hypoglossia, mandibular hypoplasia, and palatoschisis. B) E2-2 from the ERas null clone-2 all-ESC pup presented with an elongated proboscis and absence of mandibula and mouth. The pup showed alobular holoprosencephaly (middle panel), where the entire skull appeared to be occupied by neuronal brain tissue (right panel). C) The E8-1 pup, generated from ERas null ESC clone-8, lacked eyes and a mouth. The nose was enlarged and knob-like in shape, with no nostrils present. Histological examination revealed astomia with agnathia and aglossia, adontia, arrhinia, anophthalmia, alobular holoprosencephaly, and synotia. D) The E8-2 pup, derived from ESC clone-8, displayed microstoma with aglossia, mandibular hypoplasia, and palatoschisis. Additionally, the pup exhibited anophthalmia, holoprosencephaly, and the absence of the liver, spleen, most parts of the intestine, pancreas, and kidneys.

We further asked whether this phenotype was specifically associated with *ERas*-null ESCs cultured in AlbuMAX-containing medium. We switched the culture medium from 2iLA to the standard 2iL medium and cultured the *ERas*-null ESCs for at least three passages in 2iL medium before tetraploid complementation. Notably, we observed that *ERas*-null all-ESC pups (5 from ERas-2 and 3 from ERas-8) appeared phenotypically normal, devoid of the craniofacial abnormalities (data not shown). These results suggest that the craniofacial abnormalities in the *ERas*-null all-ESC pups might be linked to lipid metabolism or lipid-mediated pluripotency transitions during early embryogenesis, a topic awaits further investigation.

### Discussion

Traditional ESC culture methods, often utilizing fetal bovine serum, frequently result in issues such as heterogeneous cell populations and a reduction in pluripotency (8). Although the use of the chemical defined 2i/LIF medium has addressed some of these challenges (4), prolonged culturing of ESCs, particularly female ESCs, raises concerns related to genome integrity (9, 10, 19). This poses challenges both in the long-term maintenance of ESCs and their application in gene editing for constructing GM mouse models (9, 10, 20–22). Lipids play a pivotal role in maintaining cellular homeostasis, serving various functions such as acting as an energy source through mitochondrial fatty acid oxidation, facilitating intracellular signal transduction, and providing essential macromolecules for membrane biosynthesis during growth and proliferation (23). Our recent study has demonstrated that the addition of lipid-rich albumin, AlbuMAX, into the 2i/LIF medium can significantly improve the genomic stability and pluripotency of cultured ESCs (11). We have successfully generated all-ESC mice directly from targeted ESCs through multiple rounds of genetic manipulations in 2iLA medium. This further demonstrates the effectiveness and robustness of this medium in supporting ESC culture and genetic manipulation while maintaining pluripotency *in vitro*. Our system, which enables the generation of both male and female isogenic all-ESC mice, offers a valuable approach for studying the function of genes involved in early embryonic development and provides an alternative method for expediting the construction of GM mouse models.

We successfully obtained five ESC clones that achieved loss of function through CRISPR targeting of the *ERas* gene, as confirmed by the depletion of *ERas* protein in ESCs cultured in 2iLA medium. The validation of *ERas* gene disruption was confirmed via DNA sequencing, revealing indels in the targeted regions that caused frame shifts. Notably, three out of the five ESC clones displayed robust capabilities in generating all-ESC mice through tetraploid complementation. Remarkably, the consistent craniofacial abnormal phenotype observed among all-ESC pups generated from these different ESC clones, including the monosomic XO female all-ESC pups created by Y chromosome deletion, effectively eliminates the possibility of the phenotypic results from off-target effects. Thus, we have employed a rigorous genetic strategy and harnessed all-ESC mice generated through tetraploid complementation from ESCs and unequivocally defined the role of *ERas* in regulating craniofacial development during early embryonic stages.

*ERas* is particularly upregulated in ESCs cultured in lipid-rich 2iLA medium (11), which is associated with lipid-induced pluripotency transitions. Both *ERas*-null male and female all-ESC pups exhibit a non-Mendelian phenotype characterized by fatal craniofacial abnormalities. This remarkable phenotype can be ameliorated when ESCs are cultured in lipid-free 2iL medium. These compelling findings strongly suggest that *ERas* might play a pivotal role in craniofacial development, and this role may be intricately linked to lipids during early embryogenesis in the all-ESC pups.

### Limitations of the study

The underlying cause of this craniofacial phenotype in *ERas* null mice, which is observed exclusively in all-ESC pups and not in naturally bred pups, remains unclear. Accumulation and mobilization of lipid droplets during implantation are instrumental in the morphogenesis of the pluripotent epiblast and the formation of the pro-amniotic cavity in mouse embryos, a pivotal step in all subsequent developmental processes (24, 25). One plausible explanation is that this phenotype might be specifically linked to lipid metabolism or lipid-mediated pluripotency transitions during early embryogenesis.

Notably, this phenotype was consistently observed in all-ESC pups generated from 2iLA cultured ESCs, and it was significantly mitigated when ESCs were cultured in lipid-free 2iL medium. These findings suggest that *ERas* potentially plays a specific role in lipid metabolism, a critical process for the morphogenesis of early mouse embryos when exposed to a high-lipid environment. The loss of *ERas* function appears to disrupt this process, resulting in craniofacial defects in all-ESC pups generated from the 2iLA-ESCs. Further investigation into the mechanisms underlying this phenotype is warranted, ideally involving normal fertilized embryos.

## Methods

### Animals and oocytes

Animals were housed and prepared following the protocol approved by the IACUC of Weill Cornell Medical College (Protocol number: 2014-0061). B6D2F1 and ICR mice were obtained from Taconic Farms (Germantown, NY). Female mice, aged 6-8 weeks, were superovulated using 5 IU of PMSG (Pregnant mare serum gonadotrophin, Sigma-Aldrich, St. Louis, MO) and 5 IU of hCG (Human chorionic gonadotrophin, Sigma-Aldrich) with a 48-hour interval between injections. MII oocytes were collected from superovulated female mice 14-16 hours after hCG administration.

### Tetraploid complementation

Two-cell embryos were collected from 1.5 d ICR pregnant mice and cultured in KSOM+AA (Specialty Media) in a humidified incubator under 5% CO2 at 37 °C. The cell fusion program was carried out by an electrofusion device to produce tetraploid embryos by electrofusion. Tetraploid embryos were washed several times by KSOM+AA. Ten to fifteen EPS cells were injected into tetraploid blastocysts, and fifteen to twenty embryos were transferred to the uterus of ICR 2.5 d pseudo-pregnant recipients.

### ESC culture

ESC lines were cultured on 2i/LIF medium plus 1% AlbuMax, 2i/LIF medium containing DMEM/F12, Neurobasal, N2 and B27 supplement, glutamine, NEAA, P/S, β - mercaptoethanol and LIF (Millipore), 1 μM PD0325901, and 3 μM CHIR99021 (STEMGENT). Above mentioned reagents were all from Theromofisher unless otherwise specified.

### gRNA and CRISPR Cas9

The gRNAs were synthesized by IDT company. gRNAs were annealed to a tracrRNA (IDT) to form dual duplex gRNAs by heating at 95 °C for 5 min. Then they were incubated with Cas9 protein (IDT, HiFi Cas9 nuclease V3) for 20 min to form RNPs. The final concentration for each electroporation was 1.8 μM gRNA and 1.5 μM Cas9 nuclease.

### Electroporation of ESCs

The cells were electroporated on Neon Transfection Device (Thermo Fisher) with optimum electroporation parameters and Neon Transfection System 10 µL Kit (Thermo Fisher, Cat.No. MPK1096). Briefly, 100, 000 ES cells were washed by PBS (without Ca2+ and Mg2+) and resuspended by 10 μL cold Resuspension Buffer R. And then 0.5 μL RNP complex and 0.5 μL GFP plasmid were added. Here we used #14 (pulse voltage: 1200; pulse width: 20; pulse no.: 2) and #15 (pulse voltage: 1300; pulse width: 20; pulse no.: 2) program to perform electroporation. After electroporation, cells were seeded into 24-well containing ESC culture medium and returned to the incubator. Forty-eight hours after transfection, the GFP positive single cell was picked up by microinjection pipette and placed into 96 wells with feeder for further culturing for 4-5 days. The expanded single cell clones were passaged for further analysis.

### Genotyping

Genotyping of ESCs and mouse tails were performed by PCR using KAPA Mouse Genotyping Kit (Roche, Cat. No. KK7302). Specific protocol followed the reagent instructions. PCR was performed using specific primers under the following conditions: 95 °C for 3 min followed by 35-40 cycles of PCR (95 °C for 15 s, 60°C for 15s, and 72 °C for 120 s).

### Off-target effect analysis

Ten potential “off-target” sites of the sgRNA used in this research were given by the Guide Design Resources. PCR products of them were sequenced for off-target effect analysis. The sequences of the primers for off-target sites were listed in Table S1.

### Western blotting assays

Cells were lysed into 1× SDS loading buffer (50 mM Tris-HCl pH 6.8,5% β-mercaptoethanol, 2% SDS, 0.01% bromophenol blue, 10% glycerol) followed by sonication (Bioruptor, 2 × 30 s at high setting). Proteins were resolved on a 5–15% gradient Tris–glycine SDS–PAGE gel and semi-dry-transferred to nitrocellulose membranes. The following primary antibodies were used: ERas (Abcam, ab192868), Gapdh (Abcam, ab8245). Horseradish peroxidase (HRP)-conjugated secondary antibodies and the ECL prime western blotting system (GE Healthcare, RPN2232) were used for detection of primary antibodies. The samples were loaded on two parallel gels synchronously and then the gels were transferred to nitrocellulose membranes synchronously. Chemiluminescent signals were captured with a digital camera (Kindle Biosciences).

## References

1. Qin, Y. R., Geng, F. Q., and Wen, D. C. (2022) Generation of sex-reversed female clonal mice via CRISPR/Cas9-mediated Y chromosome deletion in male embryonic stem cells. Method Cell Biol 170, 203–210

2. Evans, M. J., and Kaufman, M. H. (1981) Establishment in Culture of Pluripotential Cells from Mouse Embryos. Nature 292, 154–156

3. Martin, G. R. (1981) Isolation of a Pluripotent Cell-Line from Early Mouse Embryos Cultured in Medium Conditioned by Teratocarcinoma Stem-Cells. P Natl Acad Sci-Biol 78, 7634–7638

4. Ying, Q. L., Wray, J., Nichols, J., Batlle-Morera, L., Doble, B., Woodgett, J., Cohen, P., and Smith, A. (2008) The ground state of embryonic stem cell self-renewal. Nature 453, 519–U515

5. Wen, D., Saiz, N., Rosenwaks, Z., Hadjantonakis, A. K., and Rafii, S. (2014) Completely ES cell-derived mice produced by tetraploid complementation using inner cell mass (ICM) deficient blastocysts. PloS one 9, e94730

6. George, S. H. L., Gertsenstein, M., Vintersten, K., Korets-Smith, E., Murphy, J., Stevens, M. E., Haigh, J. J., and Nagy, A. (2007) Developmental and adult phenotyping directly from mutant embryonic stem cells. P Natl Acad Sci USA 104, 4455–4460

7. Eggan, K., Akutsu, H., Loring, J., Jackson-Grusby, L., Klemm, M., Rideout, W. M., Yanagimachi, R., and Jaenisch, R. (2001) Hybrid vigor, fetal overgrowth, and viability of mice derived by nuclear cloning and tetraploid embryo complementation. P Natl Acad Sci USA 98, 6209–6214

8. Hackett, J. A., and Surani, M. A. (2014) Regulatory Principles of Pluripotency: From the Ground State Up. Cell Stem Cell 15, 416–430

9. Choi, J., Huebner, A. J., Clement, K., Walsh, R. M., Savol, A., Lin, K., Gu, H., Di Stefano, B., Brumbaugh, J., Kim, S. Y., Sharif, J., Rose, C. M., Mohammad, A., Odajima, J., Charron, J., Shioda, T., Gnirke, A., Gygi, S., Koseki, H., Sadreyev, R. I., Xiao, A., Meissner, A., and Hochedlinger, K. (2017) Prolonged Mek1/2 suppression impairs the developmental potential of embryonic stem cells. Nature 548, 219–223

10. Yagi, M., Kishigami, S., Tanaka, A., Semi, K., Mizutani, E., Wakayama, S., Wakayama, T., Yamamoto, T., and Yamada, Y. (2017) Derivation of ground-state female ES cells maintaining gamete-derived DNA methylation. Nature 548, 224–227

11. Zhong, L., Gordillo, M., Wang, X., Qin, Y., Huang, Y., Soshnev, A., Kumar, R., Nanjangud, G., James, D., David Allis, C., Evans, T., Carey, B., and Wen, D. (2023) Dual role of lipids for genome stability and pluripotency facilitates full potency of mouse embryonic stem cells. Protein Cell 14, 591–602

12. Takahashi, K., Mitsui, K., and Yamanaka, S. (2003) Role of ERas in promoting tumour-like properties in mouse embryonic stem cells. Nature 423, 541–545

13. Wang, X. X., Xiang, Y. L., Yu, Y., Wang, R., Zhang, Y., Xu, Q. H., Sun, H., Zhao, Z. A., Jiang, X. X., Wang, X. Q., Lu, X. K., Qin, D. D., Quan, Y. J., Zhang, J. Q., Ng, S. C., Wang, H. M., Jing, N. H., Xie, W., and Li, L. (2021) Formative pluripotent stem cells show features of epiblast cells poised for gastrulation. Cell Research 31, 526–541

14. Kinoshita, M., Barber, M., Mansfield, W., Cui, Y., Spindlow, D., Stirparo, G. G., Dietmann, S., Nichols, J., and Smith, A. (2021) Capture of Mouse and Human Stem Cells with Features of Formative Pluripotency. Cell Stem Cell 28, 2180

15. Yu, L., Wei, Y., Sun, H. X., Mahdi, A. K., Pinzon Arteaga, C. A., Sakurai, M., Schmitz, D. A., Zheng, C., Ballard, E. D., Li, J., Tanaka, N., Kohara, A., Okamura, D., Mutto, A. A., Gu, Y., Ross, P. J., and Wu, J. (2021) Derivation of Intermediate Pluripotent Stem Cells Amenable to Primordial Germ Cell Specification. Cell Stem Cell 28, 550–567 e512

16. Zhao, Z. A., Yu, Y., Ma, H. X., Wang, X. X., Lu, X. K., Zhai, Y. H., Zhang, X. X., Wang, H. B., and Li, L. (2015) The roles of ERAS during cell lineage specification of mouse early embryonic development. Open Biol 5

17. Yu, Y., Liang, D., Tian, Q., Chen, X. N., Jiang, B., Chou, B. K., Hu, P., Cheng, L. Z., Gao, P., Li, J. S., and Wang, G. (2014) Stimulation of Somatic Cell Reprogramming by ERas-Akt-FoxO1 Signaling Axis. Stem Cells 32, 349–363

18. Kwon, Y. W., Jang, S., Paek, J. S., Lee, J. W., Cho, H. J., Yang, H. M., and Kim, H. S. (2015) E-Ras improves the efficiency of reprogramming by facilitating cell cycle progression through JNK-Sp1 pathway. Stem Cell Res 15, 481–494

19. Pastor, W. A., Chen, D., Liu, W. L., Kim, R., Sahakyan, A., Lukianchikov, A., Plath, K., Jacobsen, S. E., and Clark, A. T. (2016) Naive Human Pluripotent Cells Feature a Methylation Landscape Devoid of Blastocyst or Germline Memory. Cell Stem Cell 18, 323–329

20. Buecker, C., and Wysocka, J. (2012) Enhancers as information integration hubs in development: lessons from genomics. Trends in Genetics 28, 276–284

21. Taapken, S. M., Nisler, B. S., Newton, M. A., Sampsell-Barron, T. L., Leonhard, K. A., McIntire, E. M., and Montgomery, K. D. (2011) Karotypic abnormalities in human induced pluripotent stem cells and embryonic stem cells. Nat Biotechnol 29, 313–314

22. Nguyen, H. T., Geens, M., and Spits, C. (2013) Genetic and epigenetic instability in human pluripotent stem cells. Hum Reprod Update 19, 187–205

23. Tsogtbaatar, E., Landin, C., Minter-Dykhouse, K., and Folmes, C. D. L. (2020) Energy Metabolism Regulates Stem Cell Pluripotency. Frontiers in Cell and Developmental Biology 8

24. Mau, K. H. T., Karimlou, D., Barneda, D., Brochard, V., Royer, C., Leeke, B., de Souza, R. A., Pailles, M., Percharde, M., Srinivas, S., Jouneau, A., Christian, M., and Azuara, V. (2022) Dynamic enlargement and mobilization of lipid droplets in pluripotent cells coordinate morphogenesis during mouse peri-implantation development. Nat Commun 13

25. Tatsumi, T., Takayama, K., Ishii, S., Yamamoto, A., Hara, T., Minami, N., Miyasaka, N., Kubota, T., Matsuura, A., Itakura, E., and Tsukamoto, S. (2018) Forced lipophagy reveals that lipid droplets are required for early embryonic development in mouse. Development 145

